# HIRA loss transforms FH-deficient cells

**DOI:** 10.1101/2022.06.04.492123

**Authors:** Lorea Valcarcel-Jimenez, Connor Rogerson, Cissy Yong, Christina Schmidt, Ming Yang, Victoria Harle, Victoria Offord, Kim Wong, Ariane Mora, Alyson Speed, Veronica Caraffini, Maxine Gia Binh Tran, Eamonn R. Maher, Grant D. Stewart, Sakari Vanharanta, David J. Adams, Christian Frezza

## Abstract

Fumarate Hydratase (FH) is a mitochondrial enzyme that catalyses the reversible hydration of fumarate to malate in the TCA cycle. Germline mutations of *FH* lead to HLRCC, a cancer syndrome characterised by a highly aggressive form of renal cancer(*1*). Although HLRCC tumours metastasise rapidly, FH-deficient mice develop premalignant cysts in the kidneys, rather than carcinomas (*2*). How *Fh1*-deficient cells overcome these tumour suppressive events during transformation is unknown. Here, we perform a genome-wide CRISPR/Cas9 screen to identify genes that, when ablated, enhance the proliferation of *Fh1*-deficient cells. We found that the depletion of HIRA enhances proliferation and invasion of *Fh1*-deficient cells *in vitro* and *in vivo*. Mechanistically, *Hira* loss enables the activation of MYC and its target genes, increasing nucleotide metabolism specifically in *Fh1*-deficient cells, independent of its histone chaperone activity. These results are instrumental for understanding mechanisms of tumorigenesis in HLRCC and the development of targeted treatments for patients.

## INTRODUCTION

Tumour initiation and progression requires the metabolic rewiring of cancer cells(*3, 4*). Fumarate hydratase (FH), a mitochondrial enzyme that catalyses the reversible hydration of fumarate to malate in the TCA cycle, has been identified as a *bona fide* tumour suppressor(*5*). FH loss predisposes to Hereditary Leiomyomatosis and Renal Cell Carcinoma (HLRCC), a cancer syndrome characterized by the presence of benign tumours of the skin and uterus, and a highly aggressive form of renal cancer (*1*). Its loss leads to aberrant accumulation of fumarate, an oncometabolite that drives malignant transformation(*6, 7*). Even though the link between FH loss, fumarate accumulation and HLRCC is well-known, the associated tumorigenic mechanism is it is still not fully understood(*8*). Indeed, although HLRCC tumours metastasize even when small, *Fh1*-deficient mice develop premalignant cysts in the kidneys, rather than overt carcinomas(*2*). Interestingly, these cysts are positive for the key tumour suppressor p21(*9*). Since p21 expression is a central trigger of cellular senescence, it is postulated that this process could be an obstacle for tumorigenesis in *Fh1*-deficient cells. Consistent with this hypothesis, HLRCC patients harbour the epigenetic suppression of p16, another key player of senescence. Here, we have confirmed that additional oncogenic events independent from a senescence bypass are required to allow full-blown transformation in FH deficient cells. Moreover, a genome wide CRISPR/Cas9 screen identified HIRA as a target that, when ablated, increases proliferation and invasion in *Fh1*-deficient cells. Moreover, *Fh1* and *Hira*-deficient cells lead to the development of tumours and invasive features in the kidney *in vivo*. Strikingly, *Hira* depletion in *Fh1* deficient cells controls the activation of a MYC and E2F-dependent transcriptional and metabolic program, which is known to play different oncogenic roles during tumour initiation and progression(*11, 12*). Of note, the activation of these programs is independent of H3.3 deposition into the chromatin, known to be controlled by HIRA(*13*). Overall, we have identified a novel oncogenic event occurring in FH deficient tumours, which study will be instrumental for understanding mechanisms of tumorigenesis in HLRCC and the development of targeted treatments.

## RESULTS

### *Fh1* loss impairs 2D growth and enhances migration and invasion

To investigate the oncogenic properties elicited by FH loss, we started by determining the 2D growth, cell cycle profile, and migration properties of mouse *Fh1*-proficient (*Fh1^fl/fl^), -*deficient (*Fh1^-/-CL1^* and *Fh1^-/-CL19^*), and -reconstituted (*Fh1^-/-CL1^ +pFH)* epithelial kidney cell lines we have previously generated(*14, 15*). *Fh1* loss led to a significant decrease in 2D growth, an arrest in G1 phase of the cell cycle, and a decrease in DNA synthesis determined by BrdU incorporation into DNA (**Fig.1A-C, Supplementary Fig.1A**). Of note, the overall decrease in cell proliferation was not associated with canonical senescence markers, such as β-Galactosidase activity and *Cdkn1a/Cdkn2a* gene expression (**Supplementary Fig.1B-C**), contrary to previous observations in primary epithelial cells(*9*). Using a permeable derivative of fumarate, we confirmed that fumarate accumulation alone was sufficient to induce a senescence-independent decrease in cell proliferation (**Supplementary Fig.1D-F**). In contrast to the effect on proliferation, *Fh1* loss led to increased cell migration and invasive growth of spheroids embedded in collagen I matrix (**Fig.1D-E, Supplementary Fig.1G**), consistent with the epithelial to mesenchymal transition (EMT) activation as we recently showed(*15*). Of note, despite not being senescent and presenting invasive and migratory features, *Fh1*-deficient cells were unable to form xenografts *in vivo* (**Fig.1G**). These results indicate that despite the higher invasion ability *in vitro* and the lack of senescence markers, *Fh1* loss in epithelial kidney cells is not sufficient to drive full-blown transformation. Therefore, we hypothesised that additional oncogenic events initiating transformation must occur in *Fh1*-deficient cells.

**Figure 1.**
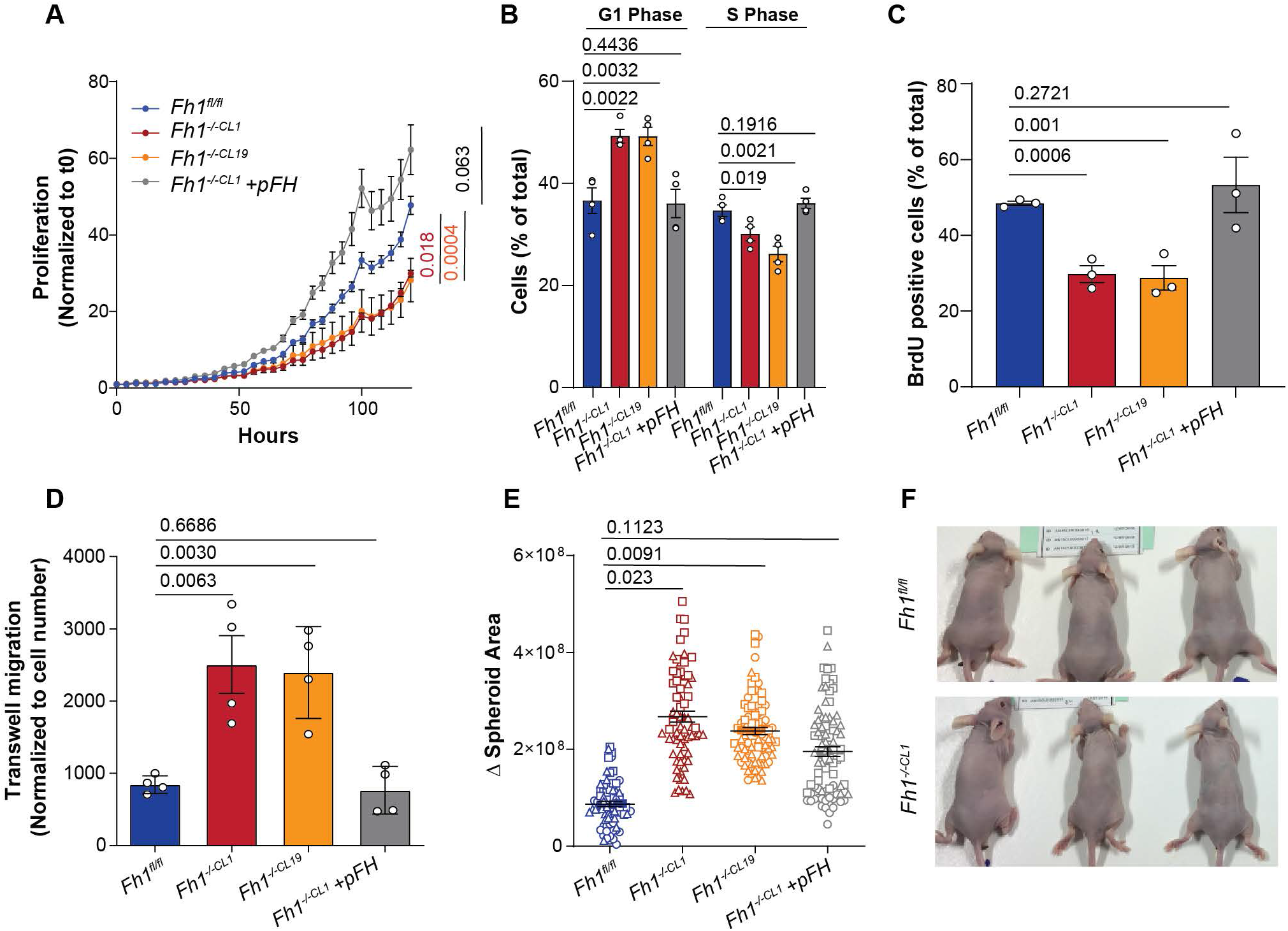
*Fh1* loss in kidney epithelial mouse cells compromises proliferation enhancing migration and invasion. **(a)** 2D growth analysis of *Fh1* proficient (*Fh1^fl/fl^),* deficient (*Fh1^-/-CL1^* and *Fh1^-/-CL19^*) and reconstituted (*Fh1^-/-CL1^ +pFH)* cell lines (n=4). Data were normalised to time 0. The last time point was used for the statistical comparison. **(b)** Cell Cycle comparisons were performed identifying differences between percentage of propidium iodide (PI) staining in G1 (physical cell growth-interphase) and S (DNA synthesis) phases of cell cycle (n=4). **(c)** DNA synthesis by means of BrdU incorporation into the DNA. Percentage of BrdU positive cells relative to total nucleus number is plotted for each condition (n=3). **(d)** Trans-well migration of cells normalised by cell number (n=4). **(e-f)** Analysis of increased spheroid area in 48 hours (n=3). At least 20 spheroids were analysed per experiment. Dots represent experiment 1, squares experiment 2 and triangles experiment 3. **(g)** Representative pictures of mice injected with *Fh1* proficient (*Fh1^fl/fl^)* and deficient (*Fh1^-/-CL1^)* cells (n=5). No tumours were visible in any of the mice. Error bars represent standard error of the mean (S.E.M). Statistic tests performed: two-tailed Student T test (a, d, e), one-tail Student t-test (b, c). Numbers represent p-value for all comparisons.

### *Hira* loss increases proliferation and invasiveness in *Fh1*-deficient cells

To identify genes that, when ablated, enhance proliferation in *Fh1-*deficient cells we performed a genome-wide CRISPR screen (**Fig.2A**) using *Fh1-* proficient (*Fh1^fl/fl^*),- deficient (*Fh1^-/-CL1^*) and -reconstituted (*Fh1^-/-CL1^ +pFH)* cells stably expressing Cas9. We transduced cells with a genome-wide guide RNA (gRNA) mouse V2 CRISPR pooled library and propagated them for 18 days, as previously described(*16, 17*). Analysis of the high-throughput sequencing data led to the identification of two significantly enriched gRNAs dependent on *Fh1* deficiency (**Fig.2B**). One of them was the metabolic enzyme *Pfkfb1*, and the other one was *Hira* (**Fig.2C**). Given the role of HIRA in senescence activation and suppression of neoplasia, we decided to investigate its role in *Fh1*-deficient cells(*18*).

**Figure 2.**
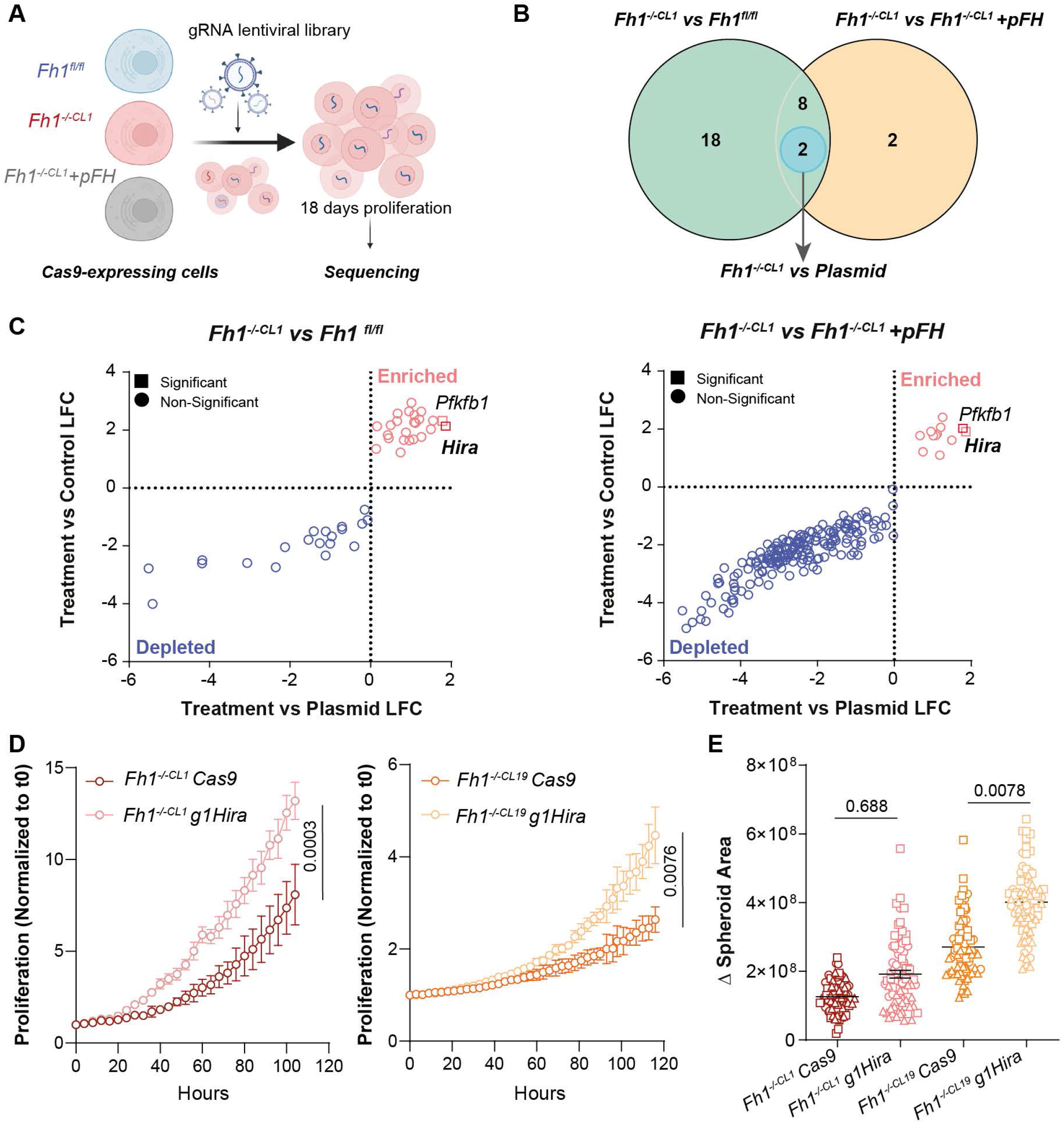
A genome-wide CRISPR/Cas9 screen identifies *Hira* loss as an oncogenic factor in *Fh1*-deficient cells. **(a)** Schematic of the CRISPR/Cas9 screen carried out. *Fh1* proficient (*Fh1^fl/fl^),* deficient (*Fh1^-/-CL1^*) and reconstituted (*Fh1^-/-CL1^ +pFH)* cell lines expressing Cas9 were transduced with a pool mouse library containing ∼90,000 gRNAs(*30*). Cells were grown for 18 days and at least 90 million cells were harvest for consequent DNA extraction and high-throughput sequencing. **(b)** Venn diagram of the comparisons performed. Two enriched gRNA dependent on *Fh1* and plasmid expression were identified (highlighted in blue). **(c)** Volcano plots showing the significantly depleted and enriched gRNAs for each comparison performed (*Fh1^-/-CL1^ vs Fh1^fl/fl^)* and (*Fh1^-/-CL1^ vs Fh1^-/-CL1^ +pFH)*. Blue and pink colours represent the comparison between *Fh1* expression conditions. Shapes refers to whether a gene is significant in the comparison treatment condition versus plasmid. This comparison gets rid of significant depleted or enriched gRNAs dependent of the corresponding gene basal expression in the cells. **(d)** 2D growth analysis of *Fh1*-deficient cells (*Fh1^-/-CL1^* and *Fh1^-/-CL19^*) under *Hira* depletion (*Fh1^-/-CL1^ g1Hira* and *Fh1^-/-CL19^g1Hira*) (n=3). Data normalised to time 0. Statistics performed comparing the values of the last time point. **(e)** Representation of the increase in spheroid area for 48 hours (n=3). At least 20 spheroids were analysed per experiment. Dots represent experiment 1, squares experiment 2 and triangles experiment 3. Error bars represent standard error of the mean (S.E.M). Statistic tests performed: two-tailed Student T test (d), one-tail Student t-test (e). Numbers represent p-value for all comparisons. LFC=Log_2_ Fold Change.

We validated the screen by generating *Hira-*deficient cells and confirmed the canonical markers of FH loss (**Supplementary Fig.3A**). Consistent with the screen results, *Hira* loss increased the 2D growth and DNA replication in two independent *Fh1*-deficient clones but had no effect in *Fh1*-proficient or reconstituted cells (**Fig.2D, Supplementary 2B**). *Hira* and *Fh1* loss were also associated with an increased percentage of cells in the S phase and BrdU incorporation into the DNA (**Supplementary Fig.2C-D**). The effect of *Hira* loss in 2D growth and cell cycle progression was also confirmed using a second gRNA (*g4Hira*) (**Supplementary Fig.2E-G**). Next, we investigated whether *Hira* loss enhanced the invasion ability of the cells. *Hira* loss further increased the invasive growth of *Fh1*-deficient cells in spheroids embedded in a collagen I matrix (**Fig.2E, Supplementary Fig.2H**). Altogether, these results indicate that the loss of *Hira* in *Fh1*-deficient cells enhances proliferation and invasion and could play a significant role in the tumorigenesis associated with FH loss.

**Figure 3.**
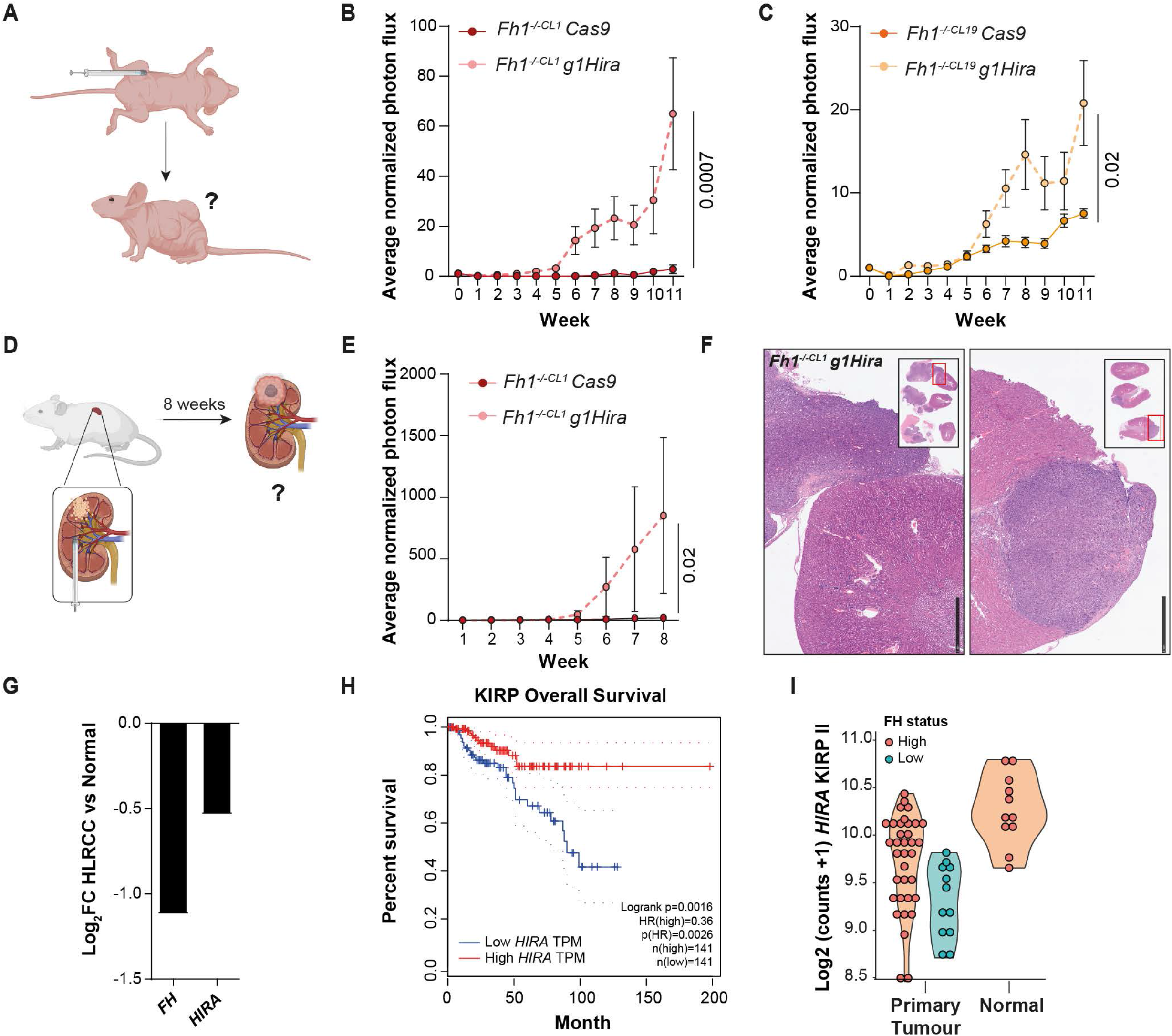
*Fh1 and Hira* deficient cells promote tumour initiation, growth and invasion *in vivo*. **(a)** Scheme of xenograft injections in the flank of nude mice. 2 million cells were injected in each flank (5 mice, 10 injections in total) and tumour initiation/growth was monitored for 11 weeks by IVIS bioluminescence imaging. **(b-c)** Xenograft tumour growth by means of Bioluminescence Imaging (BLI) and flux intensity normalised to day 0 (n=10 tumours) of *Hira* and *Fh1*-deficient cells (*Fh1^-/-CL1^ g1Hira* and *Fh1^-/-CL19^ g1Hira)*. **(d)** Scheme of orthotopic experiments carried out. Cells were injected in the kidney capsule (n=4 kidneys/condition). Tumour initiation, growth and invasion was analysed for 8 weeks by IVIS bioluminescence imaging. **(e)** Representation of luminescence signal by means of Bioluminescence Imaging (BLI) and flux intensity normalised to day 1 (day after surgery) of *Fh1*-deficient (*Fh1^-/-CL1^ Cas9)* and *Hira* and *Fh1*-deficient cells (*Fh1^-/-CL1^ g1Hira*). **(f)** Representative hematoxylin & eosin (H&A) images of the kidney injected with *Hira-* and *Fh1*-deficient cells. Tumours attached to the kidney capsule and invasive lesions within the kidney can be observed. Small square represents the whole sections of the kidney and adjacent tumours. Scale bars represent 1mm. **(g)** Analysis of FH and HIRA expression data from HLRCC patients(*21*). **(h)** Overall survival data associated to HIRA expression from Papillary type renal cancer (KIRP) using GEPIA(*22*). **(i)** Gene expression of HIRA in Papillary type II renal cancer (KIRP) comparing Normal and primary tumour samples. Tumour samples with low FH expression are represented in blue. Error bars represent standard error of the mean (S.E.M). Statistic tests performed: two-tailed Mann-Whitney U test. Numbers represent p-value for all comparisons. HLRCC: Hereditary Leiomyomatosis and renal cell cancer; TPM: Transcripts per million, FC: Fold Change.

### *Fh1 and Hira* deficient cells promote tumour initiation, growth, and invasion *in vivo*

Given the role played by *Hira* depletion in *Fh1*-deficient cells *in vitro*, we next investigated its role *in vivo*. To do so, we performed two different *in vivo* experimental approaches. First, we xenografted 1 million cells subcutaneously per condition in the flanks of nude mice and monitored tumour formation and growth for 11 weeks (**Fig.3A**). *Hira-* and *Fh1*-deficient cells significantly increased tumour initiation and formation compared to *Fh1-*deficient cells only (**Fig.3B-C, Supplementary Fig.3A**). Next, we performed orthotopic cell injection in the kidney capsule and monitored the survival and growth of the cells(*19*)(**Fig.3D**). This approach better mimics the microenvironment where FH-deficient tumours develop *in vivo*(*20*). Strikingly, tumours were detected in the kidney capsules injected with *Fh1-* and *Hira*-deficient cells, while no tumours were observed in those injected with *Fh1*-deficient cells (**Fig.3E-F, Supplementary Fig.3B-C**). Cells deficient for *Hira* and *Fh1* showed invasive potential within the kidney (**Fig.3F**), indicating a higher degree of malignancy. Of note, control cells (*Fh1^fl/fl^)* did not generate tumours *in vivo* (**Supplementary Fig.3D-E**).

To corroborate these results in human samples, we took advantage of a previously published transcriptomics analysis of 25 HLRCC patients(*21*). We confirmed the downregulation of *FH* and *HIRA* expression in HLRCC patients compared to controls (**Fig.3G**). Furthermore, we validated the downregulation of *HIRA* gene expression in kidney biopsies from two additional HLRCC patients compared to normal adjacent tissue (**Supplementary Fig.3F**).

As HLRCC predisposes to papillary type 2 renal-cell carcinoma (KIRP II), we analysed *HIRA* expression in those tumours(*1*). In addition, we used the GEPIA tool to study the association of *HIRA* expression and overall survival in kidney renal papillary cell carcinoma (KIRP)(*22*). Strikingly, *HIRA* was downregulated in primary tumours of KIRPII patients with low FH expression compared to normal tissue, and the downregulation of *HIRA* was associated with poorer overall survival in KIRP patients (**Fig.3H-I**). Together, these results confirm the previous *in vitro* results and highlight the loss of *HIRA* expression as an oncogenic event for FH deficiency *in vivo*.

### *Hira* loss in *Fh1*-deficient cells leads to a H3.3 deposition-independent MYC and E2F-transcriptional programmes activation

To gain insight into the mechanism by which *Hira* loss promotes transformation in *Fh1*-deficient cells, we performed a transcriptomic analysis followed by a gene set enrichment analysis (GSEA) of *Hira* and *Fh1*-deficient cells. This analysis showed an upregulation of EMT, MYC and E2F targets signatures (**Fig.4A**). Importantly, these transcriptional changes were genotype-specific since *Fh1* or *Hira* loss in control cells didn’t elicit the upregulation of these signatures (**Supplementary Fig.4A-B**). The activation of an EMT programme in *Fh1*-deficient cells has been previously described by our group(*15*). *Hira* loss further enhanced the downregulation of the epithelial marker E-cadherin, and the upregulation of the mesenchymal marker Vimentin by transcript and protein levels (**Supplementary Fig.4C-D**). Remarkably, when we performed the GSEA in the HLRCC patient cohort we observed MYC and E2F targets signatures as the top upregulated ones, together with a significant upregulation of the EMT signature (**Fig.4B**). Of note, the expression of these signatures was associated with poorer overall survival in KIRP patients (**Supplementary Fig.4E**).

**Figure 4.**
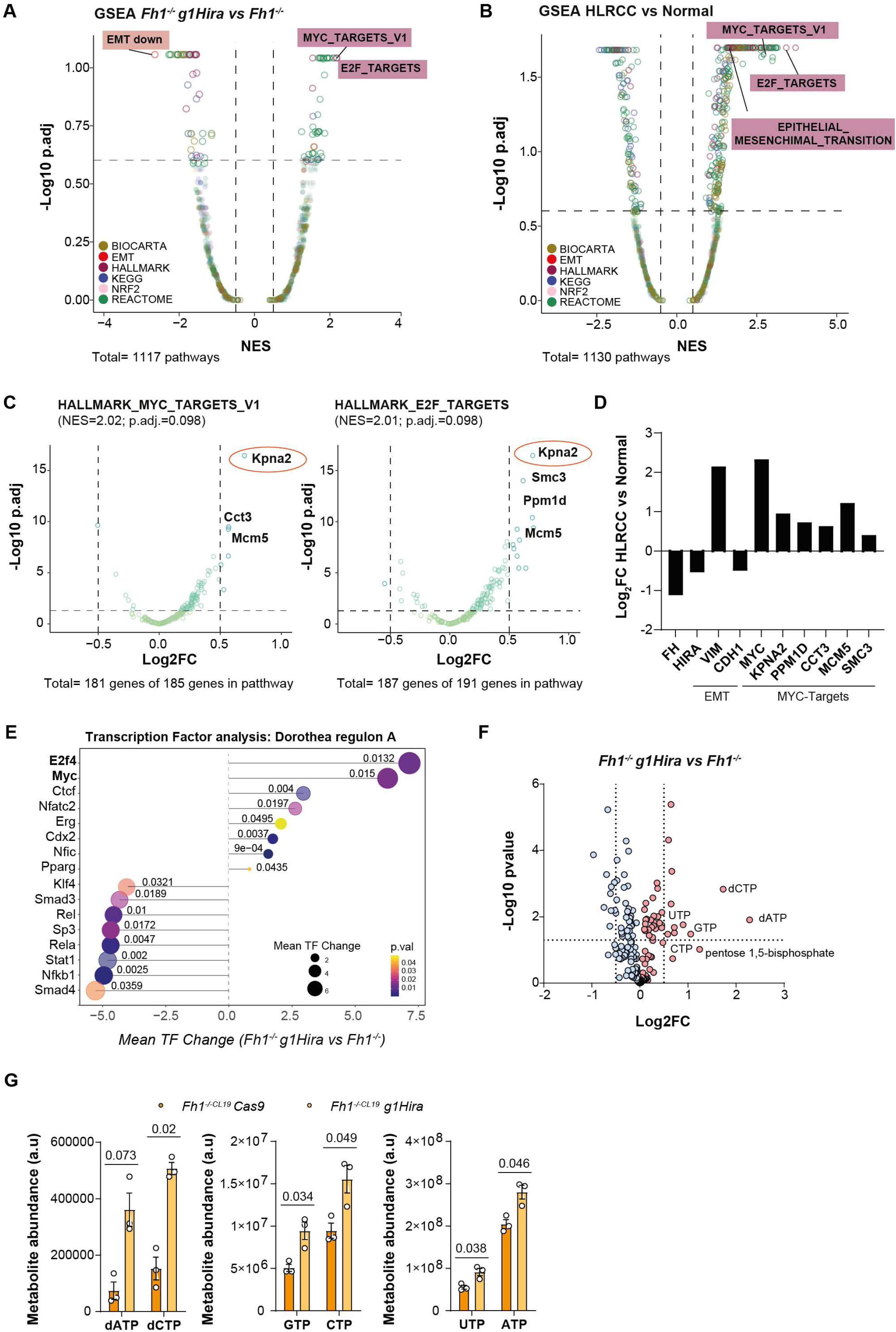
*Fh1* and *Hira* deficient cells activate an EMT programme and Myc and E2f-targets signatures. **(a)** Volcano plot representing the GSEA for *Fh1^-/-^ g1Hira vs Fh1^-/-^* Cas9 cells. Signatures are coloured depending on database represented. **(b)** Volcano plot representing the GSEA from the comparison between HLRCC and Normal patient transcriptomic data. Signatures are coloured depending on database represented. **(c)** Volcano plots of the genes present in the significantly upregulated signatures in the GSEA for *Fh1^-/-^ g1Hira vs Fh1^-/-^* Cas9 cells (MYC_Targets_V1 and E2F_Targets). **(d)** Transcriptomic expression of *FH*, *HIRA*, FH loss associated genes (*NQO1* and *HMOX1*), EMT genes (*VIM* and *CDH1*) and MYC targets (*KPNA2* and *PPM1D*) in HLRCC patient cohort. **(e)** Lollipop graph representing the mean transcription factor (TF) change in the transcriptomic data comparing *Hira* and *Fh1* deficient cells with Fh1 deficient cells alone **(f)** Volcano plot with the untargeted metabolomics performed. Nucleotides upregulated in *Fh1* and *Hira*-deficient cells. **(g)** Metabolite abundance of the nucleotides shown for Fh1-deficient (*Fh1^-/-^)* and Hira and Fh1-deficient cell lines (*Fh1^-/-^* g1Hira). NES=Normalized enrichment score. Error bars represent standard error of the mean (S.E.M). Cut off for transcriptomic volcano plots: NES +/-0.5 and p.adj=0.25 (=25%). Statistic tests performed: two-tailed Student T test (e, f). For comparisons between *Fh1*-deficient cells and *Hira* and *Fh1-*deficient cells, a paired comparison was performed. Numbers represent p-value for all comparisons.

We then investigated the most upregulated genes in MYC and E2F targets signatures in *Hira* and *Fh1*-deficient cells (**Fig. 4C**). The top upregulated factor in both signatures was the Karyopherin Subunit Alpha 2 (*Kpna2*), a nuclear transporter involved in the nucleocytoplasmic transport pathway of several tumour-associated proteins, including MYC and E2Fs(*23*). Of note, the overexpression of this transporter has been shown to promote cell proliferation (promoting the G1/S cell cycle transition), migration, invasion, and cell-matrix adhesion in several cancers, including renal cell carcinoma (RCC)(*23, 24*). Notably, a DNA replication initiation factor (MCM5) and Protein phosphatase 1D (PPM1D), a gene encoding the protein phosphatase 2c delta (PP2C*δ)* and negative regulator of cellular stress response pathways, were also upregulated as part of these signatures (**Fig.4C**)(*23, 25*). We confirmed that the downregulation of FH and HIRA was associated with the upregulation of MYC and E2F targets, as well as the EMT activation, in the cohort of HLRCC patients from Crooks D.R. et al (**Fig.4D, Supplementary Fig.5A**). Some of these transcriptional changes were validated with an additional gRNA for Hira (*g4Hira*) and in the xenograft tumours generated (**Supplementary Fig.5B-D**). Moreover, a transcription factor analysis performed revealed E2F4 and MYC as two of the top transcription factors involved in the transcriptional activation led by *Hira* loss in *Fh1*-deficient cells (**Fig4E, Supplementary Fig5E**). This validates the relevance of cell cycle progression and oncogenic transcriptional activation in the cells. Finally, we confirmed the activation of MYC-dependent pathways through an untargeted metabolomics analysis. MYC activation has been previously associated with nucleotide biosynthesis(*26*). The metabolomics analysis showed a significant upregulation of nucleotide-associated metabolites under *Hira* loss in *Fh1*-deficient cells, which validates the increased in proliferation previously shown (**Fig.4F-G, Fig.2D**).

**Figure 5.**
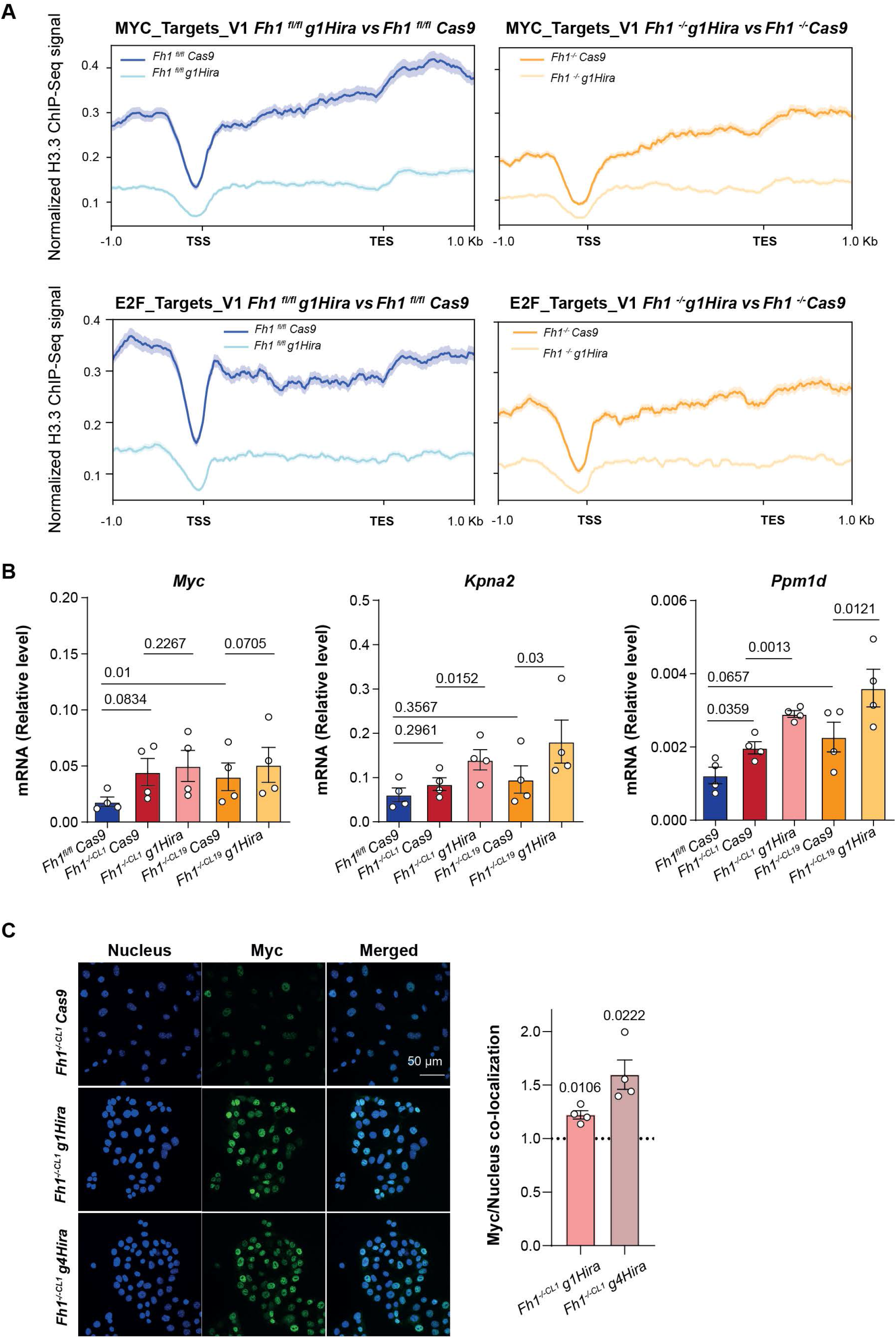
The activation of Myc and E2f-targets signatures is independent of H3.3 deposition. **(a)** Normalised ChIP-Seq signal associated with MYC and E2F target signatures expression for all conditions. Shadows represent the S.E.M. **(b)** qRT-PCR showing expression levels for *Myc*, *Kpna2* and *Ppm1d* in control (*Fh1^fl/fl^)*, *Fh1*-deficient (*Fh1^-/-CL1^*) and *Hira-* and *Fh1*-deficient cells (*Fh1^-/-CL1^ g1Hira*) (n=4). **(c)** Confocal representative images for Myc and Nuclear co-localization comparing the effect of two independent gRNAs for Hira (*g1Hira and g4Hira*) in *Fh1*-deficient cells (*Fh1^-/-CL1^*) (n=3). Dotted line represents *Fh1*-deficient cells (*Fh1^-/-CL1^ Cas9)*. β-Actin was used as a housekeeping gene. TSS=Transcription starting site, TES= Transcription end sites. Error bars represent standard error of the mean (S.E.M). Statistic tests performed: two-tailed Student T test. For comparisons between *Fh1*-deficient cells and *Hira* and *Fh1-*deficient cells, a paired comparison was performed. Numbers represent p-value for all comparisons.

We next studied how HIRA regulates the activation of MYC and E2F transcriptional signatures. HIRA, together with UBN1 and CABIN1, is part of a chaperone complex that controls the deposition of the histone variant H3.3 into the chromatin (Ray-Gallet et al., 2018). Therefore, we first assessed whether the activation of MYC, E2F and EMT – associated transcriptional programmes was due to a remodelling of H3.3 deposition in the chromatin in *Fh1* and *Hira*-deficient cells. To this aim, we performed a ChIP-Seq for H3.3 in *Fh1*-proficient and -deficient cells and analysed the effect of *Hira* loss. We observed an overall decrease in H3.3 deposition in *Hira*-deficient cells, independent of transcriptional upregulation or downregulation (**Supplementary Fig.6A**). Then, we associated the expression of MYC, E2F and EMT signatures with H3.3 normalized ChIP signal in *Fh1*-proficient and -deficient cells in the presence or absence of *Hira*. *Hira* loss resulted in a decreased H3.3 deposition into the chromatin associated to the expression of the signatures, independent of *Fh1* expression (**Fig5A, Supplementary Fig6B**). Moreover, no major differential changes were observed for H3.3 deposition in the promoter regions of the most upregulated genes (**Supplementary Fig6C**). While the activation of MYC and E2F1 target signatures is specific of *Hira* loss in Fh1-deficient cells (**Fig4A, Supplementary Fig4A-B**), the normalized H3.3 chip signal is decreased when Hira is loss in both *Fh1-*proficient and *Fh1-*deficient cells. Then, we conclude that HIRA loss may induce a transcriptional activation independent of its role controlling H3.3 deposition.

Strikingly, although gene expression analysis showed an upregulation of Myc in both *Fh1*-deficient and *Hira-* and *Fh1*-deficient cells, this was not the case for its targets *Kpna2* and *Ppm1d*, whose upregulation was specific of *Hira* loss (**Fig.5B**). Suggesting that Myc activity, rather than its expression is increased by the loss of *Hira*. Consistent with this hypothesis, we observed that MYC localisation in the nucleus was higher in *Hira* and *Fh1*-deficient cells when compared to *Fh1*-deficient cells only (**Fig.5C**). These results demonstrate that MYC transcriptional activity in *Fh1*-deficient cells may be hindered by Hira expression, compromising the activation of its transcriptional programme and that Hira loss promotes MYC binding to its transcriptional targets.

## Discussion

Although germline mutations of FH loss predispose to renal cancer in HLRCC patients, it is still unclear whether additional oncogenic events are required to transform FH-deficient cells. In this work, using a genome-wide CRISPR screening, we identified *Hira* as an oncogenic factor for *Fh1*-deficient cells *in vitro* and *in vivo*. We show that the loss of *Hira* is a prerequisite to the full activation of the proto-oncogene Myc, affecting its localisation in the nucleus and its transcriptional activity (**Supplementary Fig.7A**). Although MYC altered expression in FH-deficient cells has been hypothesised(*27*), the role of this oncogene in their transformation has not been fully investigated^17^. While HIRA has been shown to interact with c-MYC on chromatin, our work reveals an essential regulatory function of Myc activity by *Hira*, which is independent of its role as a H3.3 chaperone(*28*). Importantly, this confirms a non-canonical function of HIRA, independent of its chaperone role, as previously observed in another study(*29*). Further investigation is needed to fully assess whether this function occurs at the chromatin-level or outside of the nucleus and whether it is dependent on the deposition of the non-canonical histone H3.3 by Hira. HIRA is known to play a role in senescence, an established feature of primary FH-deficient cells triggered by fumarate. It is therefore possible that upon FH loss, the replicative arrest we observed is mediated by HIRA, at least in part by its suppressive role against Myc. Although we have not detected changes in Hira levels between FH-proficient and deficient cells, the loss of FH may affect its function on chromatin. It will be important to determine whether FH loss and the accumulation of fumarate affect the binding of Hira on chromatin, or its H3.3 deposition activity.

Overall, these results expand our understanding of how tumorigenesis occurs in *Fh1*-deficient cells and highlight the role of HIRA activating MYC-associated transcriptional programs that could lead to the development of targeted treatments.

### Limitations of the study

Although comprehensive in the study of *HIRA* loss triggering transformation in *FH*-deficient cells, this study was performed in immortalized mouse epithelial cells. Further validations would be required *in vivo*, to ascertain the effect of *HIRA* loss in a *Fh1*-deficient kidney-specific mouse model. Furthermore, it will be crucial to further investigate in the future the mechanism by which HIRA modulates MYC and E2F1-dependent transcriptional programs. In this study we hypothesize that HIRA controls the access of MYC to its transcriptional targets at chromatin level. Whether this control is directly or indirectly regulated by HIRA would be detrimental to understand its role in different tumours.

## MATERIALS AND METHODS

### Cell culture, cell lines generation and treatments

*Fh1*-proficient (*Fh1^fl/f^*), deficient (*Fh1^-/-CL1^* and *Fh1^-/-CL19^*) and reconstituted (*Fh1^-/-CL1^ +pFH*) mouse cell lines were obtained as previously described(*14, 15*). Senescence positive cells were kindly provided by Marta Pez Ribes. Cells were cultured using high glucose (4.5 g/L) DMEM (Gibco-41966-029) supplemented with 10% heat inactivated fetal bovine serum (FBS). Monomethyl-fumarate (Sigma-Aldrich) powder was resuspended in DMSO (Thermo Fischer Scientific) at 500mM and used at 400µM for 48 hours. Cells were transduced using a lentiviral packaging system with a Cas9 expressing vector (pKLV2-EF1a-Cas9Bsd, Addgene #68343) and Cas9 activity was measure using two different Cas9 reporters (pKLV2-U6gRNA5(gGFP)-PGKBFP2AGFP, Addgene # 67980 and pKLV2-U6gRNA5(Empty)-PGKBFP2AGFP, Addgene # 67979) using a LSR Fortessa (BD Bioscience) FACS. The screen was performed using the mouse improved genome-wide knockout CRISPR library V2 (pKLV2-U6gRNA5(BbsI)-PGKpuro2ABFP-W, Addgene #67988) and the different gRNA used for Hira depletion were cloned into a vector with the same backbone as the library (pKLV2-U6gRNA5(BbsI)-PGKpuro2ABFP-W, Addgene #67974). All gRNA constructs used in this study were purchased from Sigma-Aldrich. The sequences used were: g1Hira (ACATGTTTGAAACGGCCTC) and g4Hira (TAGGGAGCGGTTTCCCGCCG). The second one was designed using CRISPOR targeting exon 1 of *Hira* (http://crispor.tefor.net/). For the different Hira gRNAs, the total cell pool was used for the experiments and no single clones were selected. Luciferase expressing cells for *in vivo* experiments were generated using a Cherry-Luc construct kindly provided by Sakari Vanharanta’ s lab.

#### Lentiviral production and transduction

HEK293FT cells were transfected with the plasmid mix (plasmid of interest, PPAX2 and pMD2.G) using Lipofectamine 3000 (Thermo Fisher) as a transfection reagent diluted in Opti-MEM (Thermo Fischer). The media containing the virus was collected 48-72 hours post-transfection and filtered using a 0.45µM sterile filter. Cells were transduced twice with the lentiviral supernatant in the presence of 8µg/mL of polybrene (Millipore). Selection of cells was performed 24 post/transduction using Puromycin (2µg/ mL) and Blasticidin (10 µg/ mL) both from Gibco/Thermo Scientific.

### Animals

All animal experiments were performed in accordance with protocols approved by the Home Office (UK) and the University of Cambridge ethics committee (PPL PFCB122AA). For xenograft subcutaneous injection, 7 weeks old female NOD/SCID mice obtained from Charles River Laboratories were injected in each flank with one million cells diluted 1:1 in matrigel:PBS. At the experimental endpoint tumours, if existing, were snap frozen for further molecular analysis. For orthotopic experiments, at least 4 weeks old NSG mice obtained from Charles River Laboratories were used and the experiment was carried out as previously described(*19*). Briefly, two million cells diluted 1:3 in matrigel:DMEM were injected in the kidney capsule. Tumor initiation and growth was monitored by IVIS bioluminescence imaging (PerkinElmer) in both *in vivo* experiments.

#### Immuno-histochemistry staining (IHC)

Kidneys were collected and fixed overnight with neutral formalin 4% and washed with PBS and 70% ethanol for 15 minutes. Lungs were embedded in paraffin, sectioned, and stained with H&E by the human research tissue bank and histopathology research support group from the Cambridge University Hospitals-NHS Foundation. Different kidney sections were imaged using Slidescanner microscope (Hamamatsu S360).

### Patient samples

Patient samples were obtained by Dr. Maxine GB Tran at the Royal Free Hospital (RFH) or Prof. Eamonn R. Maher upon informed consent for genetic studies and with evaluation and approval from the corresponding ethics committees in accordance with the Declaration of Helsinki (NHS REC 16-WS-0039, South Birmingham Research Ethics Committee).

### Genome-wide CRISPR-Cas9 screen and data analysis

Cas9 expressing cells were generated and the lentiviral gRNA library was produced using HEK293FT cells as described above. A total of 150 million cells were transduced with the pooled library at a low MOI (<0.3) to ensure that >85% of cells had a single gRNA integration, resulting in at least 500x gRNA representation. Media containing the lentiviral particles was removed from the cells the following day and puromycin selection was applied for the following three days. Cells were then grown for 18 days and at least 90 million cells were harvested at the end of the assay for genomic DNA extraction. DNA was extracted using the QIAamp DNA mini kit (Qiagen) and amplified for the region containing the gRNAs, followed by high-throughput, 19bp single end sequencing (Illumina-C HiSeq 2500).

Guides were quantified against the Yusa Mouse V2 library (Addgene #67988) using crisprReadCounts v1.3.1 (https://github.com/cancerit/crisprReadCounts). Raw count normalisation to plasmid and copy number correction was performed using pyCRISPRcleanR version 2.0.8 (https://github.com/cancerit/pyCRISPRcleanR). The corrected counts were used as inputs for pairwise comparisons with MAGeCK version 0.5.9.2 to identify significantly enriched and depleted guides or genes using ‘mageck test’ with normalisation disabled (‘--norm-method non’) (https://sourceforge.net/projects/mageck) (*31*). Quality control and post-analysis tables and plots were generated in RStudio v1.2.1578 (R version 3.6.1). The detailed R scripts for the screen analysis are available on GitHub (https://github.com/team113sanger/Fumarate_Hydratase_FH_CRISPR).

### Cellular and molecular assays

#### Immunofluorescence

Immunofluorescence was performed as previously described(*32*). Briefly, cells were fixed with 4% paraformaldehyde (PFA) in PBS for 15 min, then washed 3 times with PBS. Cells were permeabilised with 0.3% Triton X-100 in 4% BSA for 20 min, followed by 3 PBS washes. Cells were then blocked with 4% BSA for 30 minutes, followed by incubation with primary antibodies in 4% BSA overnight at 4°C. After 5 washes with PBS, cells were incubated with appropriate secondary antibodies (1:1000) for 2 h at RT. After 3 washes in PBS, coverslips were mounted onto slides using the Pro Long Gold antifade mountant (Thermo Fischer). For Phalloidin staining, cells were incubated with 1:500 diluted 488 Phalloidin (Thermo Fischer) and washed 3 times with PBS. Images were taken using either the SP5 or Stellaris confocal microscopes (Leica).

#### Invasive growth and migration trans-well assays

The invasive growth assay was performed as previously described(*32*). Briefly, cells (1000 cells/drop) were maintained in drops (25 µL/drop) with DMEM and 20% methylcellulose (Sigma M0387) on the cover of a 100-mm culture plate. Drops were incubated at 37°C and 5% CO2 for 72 hours. Once formed, spheroids were collected, resuspended in collagen I solution (Advanced BioMatrix PureCol), and added to 24-well plates. After 4 hours, DMEM medium was then added on top of the well and to calculate the increased spheroid area pictures were taken using a EVOS microscope (Thermo Fischer) at days 0 and 2. For invasive growth quantification, an increase in the area between day 0 and day 2 was calculated using FiJi software. The migration trans-well assay was performed as previously described(*32*). In brief, chambers with membranes of 8 μm pores (BD Falcon) were used. Cells (50,000 cells/well) were re-suspended in 0.1 % FBS DMEM and seeded in the upper part of the chamber. In the bottom part of the well, 1.4mL of complete DMEM were added. Plates were maintained at 37°C and 10% CO2 for 24 hours. Migration was stopped washing the wells twice with PBS and using a cotton bud to remove the remaining cells of the upper part of the membrane, being careful not to compromise it. The membrane was fixed with 10% formalin (15 minutes at 4 °C) and stained with crystal violet. Cells were counted under the EVOS microscope (Thermo Fischer).

#### Cell growth, BrdU incorporation, β-Galactosidase assay and cell cycle analysis

Cell proliferation was analysed using the Incucyte SX5 by means of phase-contrast sharpness for 4-6 days or through crystal violet staining as previously described(*33*). Briefly, 5,000 cells were plated onto 24-well plates (at least 3 replicates/experimental conditions for each cell line) and at each time point cells were washed with PBS and fixed with 4% buffered formalin. Once all the time points were collected, the cells were washed with PBS and incubated with 0.1% crystal violet diluted in 20% methanol. Once the cells were stained the plates were washed with water and dried overnight. To quantify the staining differences, cells were diluted in 0.5 mL 10% acetic acid for 30 minutes at RT and quantified using TECAN spectrophotometer reading the absorbance at 595 nm. The senescence assay was performed using the senescence β-Galactosidase staining kit from Cell Signaling Technology (#9860) following manufacturer instructions. BrdU staining was performed as previously described(*34*). For BrdU incorporation, cells were seeded on coverslips in 12-well plates and after 2 days, cells were incubated with 3 μg/mL BrdU (Sigma). Cells were fixed with 4% paraformaldehyde, permeabilised with 1% Triton X-100 and incubated with a monoclonal anti-BrdU antibody (ab6326) at a 1:100 dilution. Images were obtained using a SP5 confocal microscope. At least three different areas per coverslip were quantified. Cell cycle analysis was performed using propidium iodide (PI) staining. 200,000 cells/well were seeded in 6-well plates and grown for 2 days. Cells were collected, resuspended in 1mL PBS and fixed while vortexing adding drop by drop 2.5 mL of cold absolute ethanol. Cells were stored at -20°C overnight. The next day, samples were centrifuged and washed once with PBS. Cell pellets were then resuspended in propidium iodide 1µg/mL (ab14083) solution with 0.05% triton X-100 and 25µg/mL RNAse (Thermo Fisher 12091021). Samples were analysed using a LSR Fortessa (BD Bioscience) FACS.

### RNA extraction and transcriptomic analysis

For RNA assays, 300,000 cells were plated onto a 6-well plate. The day after, cells were washed in PBS and then RNA was extracted using RNeasy kit (Qiagen) following the manufacturer’s protocol. RNA was eluted in water and then quantified using Nanodrop (Thermo Fisher). 1µg of RNA was reverse transcribed using the Quantitect Reverse Transcription kit (Thermo Fisher). For real-time qPCR, cDNA was run using Taqman assay primers (Thermo Scientific) and Taqman Fast 2X master mix (Thermo Scientific). β-Actin (ACTB) was used as the endogenous control for *in vitro* and *in vivo* experiments and RPLP0 for patient samples. The different biological replicates and gene expression differences were analysed using the ΔCt formula. For tissue samples, a maximum of 30 mg/tissue were homogenised using the Precellys® tissue homogeniser. The RNA extraction was performed with the RNeasy kit (Qiagen) following manufacturer’s instructions and gene expression analysis was performed as previously mentioned. The references of primers used can are in **Supplementary Table 1**. For RNA-Seq samples preparation, RNA was extracted as mentioned before and further purified using the RNA Clean and Concentrator kit (Zymo Research). The RNA-Seq was done on a single end run using the Truseq Stranded mRNA from Illumina, and the library preparation was performed following the manufacturer’s instructions. The sequencing was done on a 75-cycle high output NextSeq 500 kit. The differential gene expression analysis was done using the counted reads and the R package edgeR version 3.26.5 (R version 3.6.1)(*35*). Prior to running EdgeR genes were filtered that didn’t have at least 5 cpm in at least half of the samples. Pairwise comparisons were run for Cl19 vs Cl19_gHira, Cl19 VS Fl, Fl VS Fl_gHira using the exact test and adjusted for multiple testing using Benjamini-Hochberg.

### Protein lysates and western blot

300,000 cells/well were seeded in 6-well plates. The day after, cells were washed in PBS and then lysed on ice with 80µL/well RIPA buffer (150mM NaCl, 1%NP-40, Sodium deoxycholate (DOC) 0.5% sodium dodecyl phosphate (SDS) 0.1%, 25mM Tris) supplemented with protease and phosphatase inhibitors (Protease inhibitor cocktail, Phosphatase inhibitor cocktail 2/3, Sigma Aldrich) for 5 minutes. Cells extracts were scraped and further lysed in a roller for 15 minutes. Protein quantification was done using a BCA kit (Pierce) following the manufacturer’s instructions. Absorbance was read using TECAN spectrophotometer at 562 nm. Samples were resuspended in the Bolt Loading buffer 1X (Thermo Scientific) and 10-20 µg of protein were loaded into 4-12% Bis-Tris Bolt gel and run at 150-200V constant for 1h in Bolt MOPS/MES 1X running buffer (Thermo Scientific). Dry transfer of the proteins to a nitrocellulose membrane was done using IBLOT2 (Thermo Scientific) for 12 minutes at 20V. Membranes were incubated in 5% non-fat milk diluted in TBS 1X +0.01 % Tween-20 (TBS-T) for 30 minutes. Primary antibodies were incubated ON at 4°C. Calnexin antibody was purchased from Abcam (ab22595), Fh1 antibody was purchased from Origene (TA500681), Myc antibody from Cell Signaling Technologies (18583S), Cdh1 from BD (610181) and Hira WC119 clone from Merck (04–1488). The day after, membranes were washed three times in TBS-T 1X and then secondary antibodies (conjugated with 680 or 800 nm fluorophores, Li-Cor) incubated for 1h at room temperature at 1:5000 dilution in 5% non-fat milk diluted in TBS-1. T. Images were acquired using Image Studio lite 5.2 (Li-Cor) on Odyssey CLx instrument 875 (Li-Cor).

### ChIP-Seq

2 × 10^7^ cells were crosslinked in 1% formaldehyde for 10 minutes at room temperature and quenched with 0.125M glycine for at least 5 minutes. Cells were washed twice with 1 x PBS and scraped into ice-cold 1 x PBS supplemented with protease inhibitor cocktail. Cells were snap frozen at this point and stored at – 80°C until further use. Protein A Dynabeads were blocked and incubated with 2 µg anti-H3.3 antibody and 1 µg Spike-in antibody. Cells were lysed in serial rounds of lysis buffer 1, lysis buffer 2 and lysis buffer 3. Chromatin was sonicated for 10-15 cycles using a Diagenode Pico, and then supplemented with 20 ng Spike-In Drosophila chromatin. Chromatin was then incubated with beads overnight. Beads were washed five times with RIPA buffer and washed with a final 1 x TE wash. DNA was eluted in elution buffer and purified using a DNA clean and concentrator kit. Eluted DNA was quantified using a QuBit. DNA libraries were prepared using TruSeq ChIP sample prep kit (Illumina) and sequenced on a NextSeq 550 (Illumina) platform.

For data analysis, sequencing reads were aligned to mm10 and dm6 using Bowtie2 v2.2.3(*36*). Reads aligning to the Drosophila genome were counted and used to generate scale factors. BAM files were then scaled to the sample with the lowest number of Drosophila reads. Only reads with a mapping quality >q30 were retained. Replicates were merged and peak calling was performed using MACS2 v2.1.1(*37*) using default parameters with additional –SPMR parameter. bedGraph files were converted to bigwig using BedGraphtoBigWig script and visualised in the IGV Genome Browser.

Metagene tag density plots were generated using computeMatrix and plotProfile tools from the deepTools package(*38*). Correlation of biological replicates was visualised using multiBigwigSummary and plotCorrelation from the deepTools package.

### Metabolomics

HILIC chromatographic separation of metabolites was achieved using a Millipore Sequant ZIC-pHILIC analytical column (5 µm, 2.1 × 150 mm) equipped with a 2.1 × 20 mm guard column (both 5 mm particle size) with a binary solvent system. Solvent A was 20 mM ammonium carbonate, 0.05% ammonium hydroxide; Solvent B was acetonitrile. The column oven and autosampler tray were held at 40 °C and 4 °C, respectively. The chromatographic gradient was run at a flow rate of 0.200 mL/min as follows: 0–2 min: 80% B; 2-17 min: linear gradient from 80% B to 20% B; 17-17.1 min: linear gradient from 20% B to 80% B; 17.1-22.5 min: hold at 80% B. Samples were randomized and analysed with LC–MS in a blinded manner with an injection volume was 5 µl. Pooled samples were generated from an equal mixture of all individual samples and analysed interspersed at regular intervals within sample sequence as a quality control.

Metabolites were measured with a Thermo Scientific Q Exactive Hybrid Quadrupole-Orbitrap Mass spectrometer (HRMS) coupled to a Dionex Ultimate 3000 UHPLC. The mass spectrometer was operated in full-scan, polarity-switching mode, with the spray voltage set to +4.5 kV/-3.5 kV, the heated capillary held at 320 °C, and the auxiliary gas heater held at 280 °C. The sheath gas flow was set to 55 units, the auxiliary gas flow was set to 15 units, and the sweep gas flow was set to 0 unit. HRMS data acquisition was performed in a range of *m/z* = 70–900, with the resolution set at 70,000, the AGC target at 1 × 10^6^, and the maximum injection time (Max IT) at 120 ms. Metabolite identities were confirmed using two parameters: (1) precursor ion m/z was matched within 5 ppm of theoretical mass predicted by the chemical formula; (2) the retention time of metabolites was within 5% of the retention time of a purified standard run with the same chromatographic method. Chromatogram review and peak area integration were performed using the Thermo Fisher software Tracefinder 5.0 and the peak area for each detected metabolite was normalized against the total ion count (TIC) of that sample to correct any variations introduced from sample handling through instrument analysis. The normalized areas were used as variables for further statistical data analysis.

### Statistical analysis and coding

n values represent the number of independent experiments performed or the number of individual mice. For each independent *in vitro* experiment, at least three biological replicates were used (except from western blots where technical replicates are represented) and a minimum number of three experiments were done to ensure adequate statistical power. For *in vitro* experiments, Student T test was applied for two component comparisons. For *in vivo* experiments, non-parametric Mann-Whitney exact test was used. Statistical analyses involving fold changes were analysed using the one-sample T test with a null hypothesis of 1. Two-tail t-test statistical analysis was used when testing for differences between two conditions. Error bars displayed on graphs represent the mean -/+ standard error of the mean (S.E.M). All statistical analyses were performed using GraphPad PRISM 9 software. Figures were compiled using either Adobe Illustrator or BioRender. The KIRP patient survival analysis was performed using GEPIA using the overall survival and median cutoff of 50% (http://gepia.cancer-pku.cn/index.html)(*22*). KIRP II patient gene expression counts were downloaded from TCGA based on patient identifiers from Chen F. et al (*39*). Samples were stratified based on FH expression into two groups (low=bottom 25% (log2(counts + 1) <= 11.8), high=>25% (log2(counts + 1) > 11.8)). Code and data are available at https://github.com/ArianeMora/KIRP_PE2.

Gene set enrichment analysis (GSEA) was performed by recapitulating on the gene sets published on the Molecular Signatures Database (MsigDB) using the packages fgsea (v1.20.0) and GSEABase (v1.56.0)(*40–42*). The EMT genset was generated by manually curating the gene list published by Taube et al(*43*). All gene names were translated into mouse gene names prior to the analysis using scibiomart (v. 1.0.2), a wrapper around the API from BioMart. Plots were generated using the EnhancedVolcano package (v.1.12.0) (https://github.com/kevinblighe/EnhancedVolcano). Detailed code can be found under https://github.com/ChristinaSchmidt1/Oncogenic_events_in_HLRCC. RNASeq data from HLRCC patients’ primary tumours and paired adjacent tissue were downloaded from GEO database (GSE157256) analysed as described under https://github.com/ChristinaSchmidt1/Oncogenic_events_in_HLRCC. In brief, differential expression analysis have been performed using DESeq2 (v1.34.0)(*44*) and genes set enrichment analysis (GSEA) was performed as previously described(*40–42*). The EMT geneset was generated as described before(*43*). Plots were generated using the EnhancedVolcano package (v1.12.0) (https://github.com/kevinblighe/EnhancedVolcano).

For the transcription factor (TF) analysis we used the normalised count data from the EdgeR (for details see RNAseq analysis) as the input. Moreover, we use dorothea https://bioconductor.org/packages/release/data/experiment/html/dorothea.html) (v.1.6.0), a mouse TF regulon collection, filtering for confidence = “A”(*45, 46*). DoRothEA is a gene regulatory network containing signed transcription factor (TF) – target gene interactions. DoRothEA regulons, the collection of a TF and its transcriptional targets, were curated and collected from different types of evidence for both human and mouse. A confidence level was assigned to each TF-target interaction based on the number of supporting evidence. The TF analysis is performed using dorothea’s function “run_viper”, which is based on the viper package (v. 1.28.0)(*47*). To compare the conditions of interest, we calculated the mean of the biological replicates and calculated the TF change by subtracting one condition from the other. The p-value was calculated using the t-test. Detailed code information can be found in https://github.com/ChristinaSchmidt1/Oncogenic_events_in_HLRCC. All data are available in the manuscript or the Supplementary materials. All materials generated in this study are available from all the authors.

## ACKNOWLEDGEMENTS

We thank Dr Alexandria Karcanias and Dr Julien Bauer (Cambridge Genomic Services, Department of Pathology, University of Cambridge) for the RNA-seq library preparation and sequencing and the human research tissue bank and histopathology research support group from the Cambridge University Hospitals-NHS Foundation for the kidney tissue processing. We thank Jianfeng Ge for providing us with the Cherry-Luciferase expressing construct. We thank the Frezza lab for their valuable input on the project. We thank the MRC Cancer Unit facilities and kitchen media.

C.F is supported by the Medical Research Council (MRC_MC_UU_12022/6), the CRUK Programme Foundation award (C51061/A27453), ERC Consolidator Grant (ONCOFUM, ERC819920) and by the Alexander von Humboldt Foundation in the framework of the Alexander von Humboldt Professorship endowed by the Federal Ministry of Education and Research.

L V-J was supported by the long-term FEBS fellowship and by the CRUK Programme Foundation award to C.F. (C51061/A27453).

C.S work was funded by the European Union’s Horizon 2020 research and innovation programme under the Marie Skłodowska-Curie grant agreement No 722605., and the Alexander von Humboldt Professorship to C.F.

C.R. was supported by the ERC Consolidator Grant (ONCOFUM, ERC819920) to C.F.

C.Y. is funded by the Wellcome Trust (RG92001) and The Urology Foundation (G100328).

G.D.S. and E.R.M. is supported by the Cancer Research UK Cambridge Centre [C9685/A25177] and NIHR Cambridge Biomedical Research Centre (BRC-1215-20014). The views expressed are those of the author(s) and not necessarily those of the NIHR or the Department of Health and Social Care. The University of Cambridge has received salary support (ERM) from the NHS in the East of England through the Clinical Academic Reserve.

S.V was supported by Medical Research Council (MC_UU_12022/7).

DJA, VH and VO are supported by the Welcome Trust (WT206194), ERC (319661) and Cancer Research UK (C20510/A21717).

## AUTHOR CONTRIBUTIONS

Conceptualization: LVJ, CF.

Methodology: LVJ, CY, AS, VC, GS, SV, CR, MY, VH.

Investigation: LVJ, CF.

Analysis: CS, VO, DA, AM.

Visualization: CS, AM.

Patient samples: MGBT, ERM.

Writing-original draft: LVJ.

Writing-review and editing: CF with the support of all the authors.

## DATA AND MATERIALS AVAILABILITY

Metabolomics, RNA-seq and ChIP-seq data have been deposited and will be publicly available as of the date of publication. Accession numbers will be included in the methods section. All original code has been deposited at GitHub and will be publicly available as of the date of publication. The coding information is also available in the methods section. Any additional information required to reanalyse the data reported in this paper is available from the lead contact upon request.

---

**Supplementary Figure 1.**
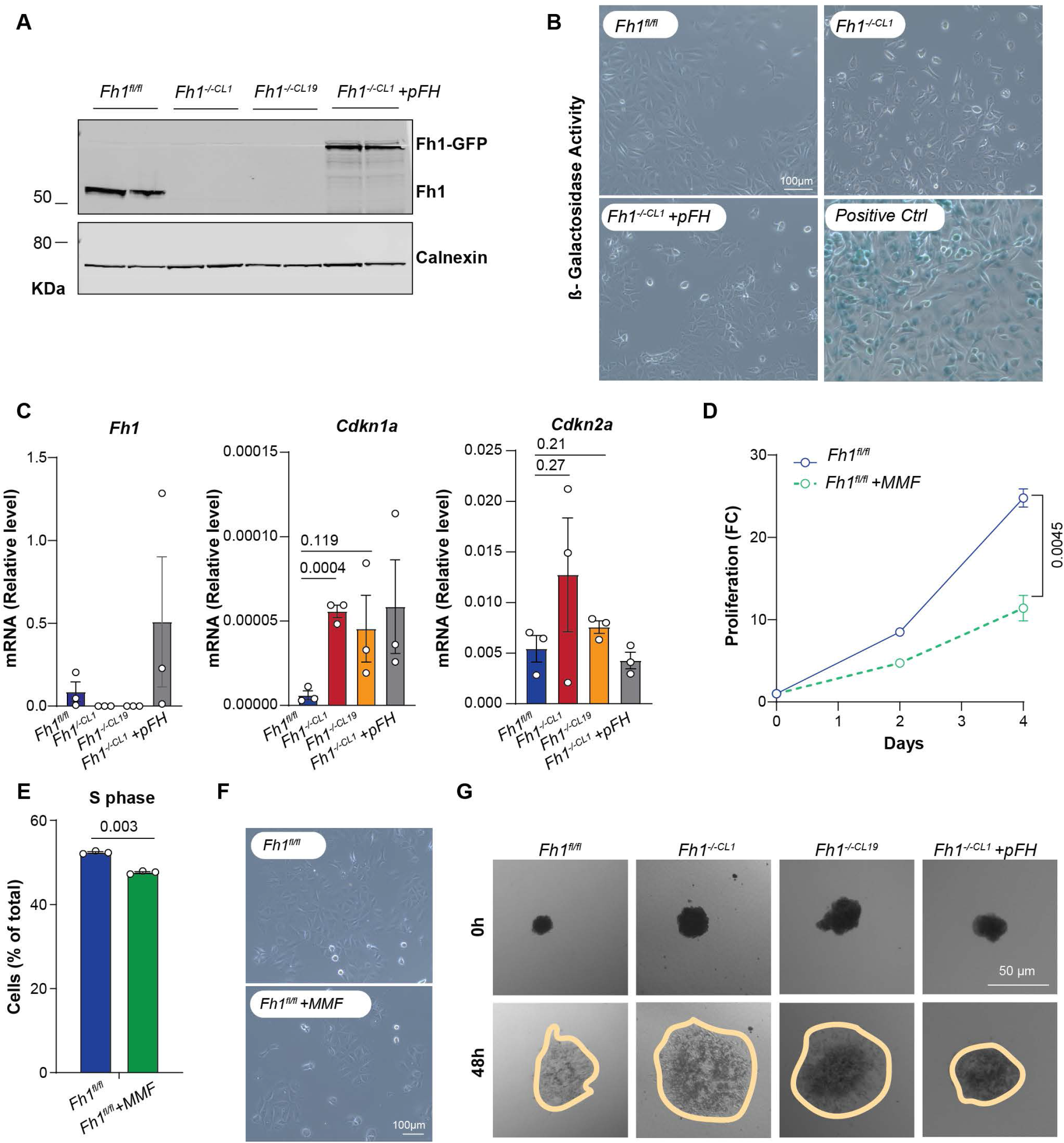
**a)** Immunoblot of Fh1 expression in *Fh1* proficient (*Fh1^fl/fl^),* deficient (*Fh1^-/-CL1^* and *Fh1^-/-CL19^*) and reconstituted (*Fh1^-/-CL1^ +pFH)* cell lines (One representative blot shown out of >5). **(b)** Representation of β-Galactosidase activity assay images for all cell lines (one representative experiment shown out of 3). No blue (β-Galactosidase activity-positive) cells were observed in comparison with a positive senescent cell line (A375 cells treated with Cisplatin). **(c)** qRT-PCR showing expression levels for *Fh1*, *Cdkn1a* (p21) and *Cdkn2a* (p16) (n=3). β-Actin was used as a housekeeping gene. **(d)** 2D growth of control cells (*Fh1^fl/fl^)* with/without monomethylfumarate (MMF) treatment (400µM). Values normalized to day 0 (n=3). Statistics performed comparing the values of the last time point. **(e)** Cell cycle analysis by means of percentage of PI staining in the S phase of the cell cycle of cells treated with MMF (n=3). **(f)** Representative images of β-Galactosidase activity in cells treated with MMF (one experiment shown out of 3). No blue (β-Galactosidase activity-positive) cells were observed in comparison with a positive senescent cell line (data not shown). **(g)** Representative images of spheroids included in collagen I matrix at 0 and 48 hours from *Fh1* proficient (*Fh1^fl/fl^),* deficient (*Fh1^-/-CL1^* and *Fh1^-/-CL19^*) and reconstituted (*Fh1^-/-CL1^ +pFH)* cell lines. One representative experiment shown out of 3. Yellow lines highlight the spheroid area. Error bars represent standard error of the mean (S.E.M). Statistic tests performed: two-tailed Student T test (c), one-tail Student t-test (d, e). Numbers represent p-value for all comparisons.

**Supplementary Figure 2.**
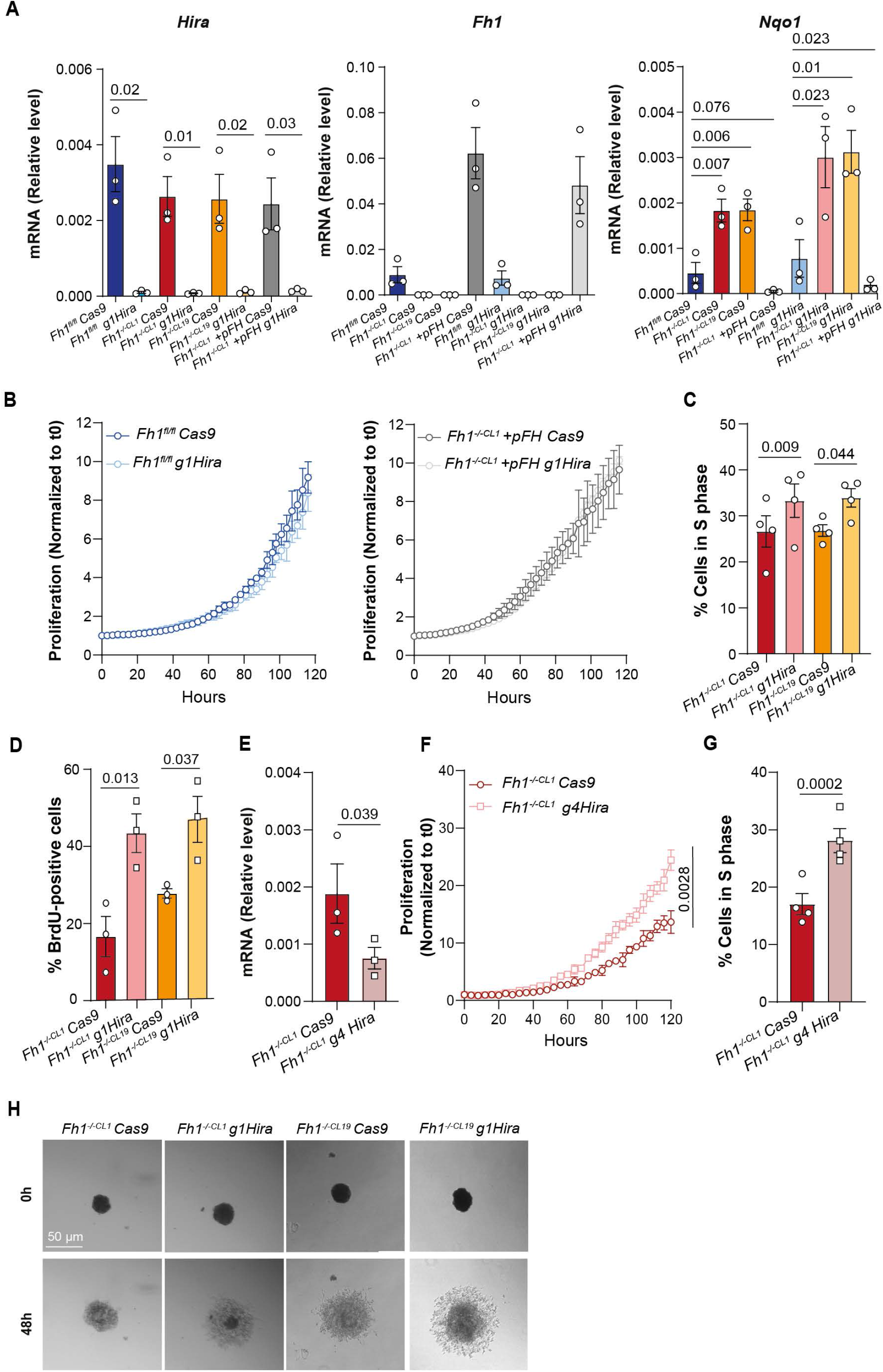
**(a)** qRT-PCR showing expression levels for *Hira*, *Fh1* and *Nqo1* (n=3) in *Fh1* proficient (*Fh1^fl/fl^ Cas9),* deficient (*Fh1^-/-CL1^ Cas9* and *Fh1^-/-CL19^ Cas9*) and reconstituted (*Fh1^-/-CL1^ +pFH Cas9)* cells alone or under *Hira* depletion (*g1Hira*) (n=3). **(b)** 2D growth analysis of *Fh1* proficient (*Fh1^fl/fl^)* and reconstituted (*Fh1^-/-CL1^ +pFH)* cell lines alone or under *Hira* loss (*g1Hira*) (n=3). Data normalised to time 0. Statistics performed comparing the values of the last time point. **(c)** Cell cycle analysis by means of percentage of cells in the S phase of cell cycle of *Fh1*-deficient cells (*Fh1^-/-CL1^ Cas9* and *Fh1^-/-CL19^ Cas9*) alone or under *Hira* deficiency (*g1Hira*). **(d)** DNA synthesis analysis by means of % of BrdU incorporation in *Fh1*-deficient cells (*Fh1^-/-CL1^* and *Fh1^-/-CL19^*) and *Hira* and *Fh1*-deficient cells (*Fh1^-/-CL1^ g1Hira* and *Fh1^-/-CL19^g1Hira*) (n=3). (**e**) qRT-PCR showing expression levels for *Hira* in Fh1-deficient cells (*Fh1^-/-CL1^ Cas9)* with a second gRNA (*g4Hira*). **(f)** 2D growth analysis of *Fh1*-deficient cells (*Fh1^-/-CL1^ Cas9)* and *Hira* and *Fh1*-deficient cells (*Fh1^-/-CL1^ g4Hira).* Values normalized to day 0 (n=3). Statistics performed comparing the values of the last time point. **(g)** Effect of Hira loss with a second gRNA (g4Hira) in *Fh1*-deficient cells (*Fh1^-/-CL1^ Cas9)* on the cell cycle progression by means of percentage of cells in the S phase (n=4). (**h**) Representative images of spheroids included in collagen I matrix at 0 and 48 hours from *Fh1*-deficient cells (*Fh1^-/-CL1^ Cas9, Fh1^-/-CL19^ Cas9)* and *Hira-* and *Fh1*-deficient cells (*Fh1^-/-CL1^ g1Hira, Fh1^-/-CL19^ g1Hira)* (n=3). Error bars represent standard error of the mean (S.E.M). Statistic tests performed: one-tailed Student T test. Numbers represent p-value for all comparisons.

**Supplementary Figure 3.**
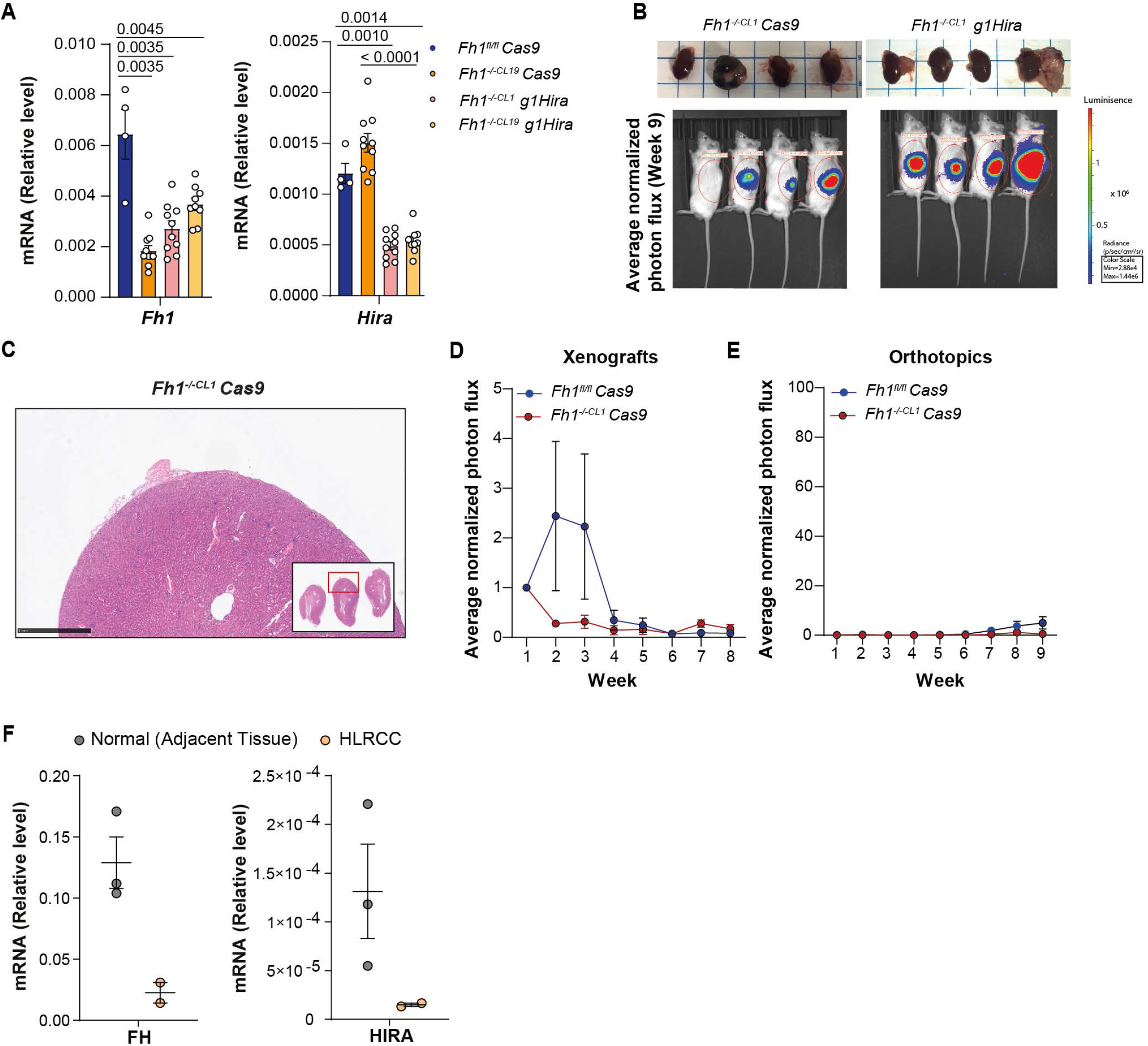
**(a**) qRT-PCR showing expression levels for *Fh1* and *Hira* in tumour xenografts from control (*Fh1^fl/fl^ Cas9)*, *Fh1*-deficient (*Fh1^-/-CL19^ Cas9)* and *Hira-* and *Fh1*-deficient cells (*Fh1^-/-CL1^ g1Hira* and *Fh1^-/-CL19^ g1Hira)* (n=10 for all conditions except for *Fh1^fl/fl^* n=4). β-Actin was used as a housekeeping gene. **(b)** Images from kidneys injected with *Fh1*-deficient (*Fh1^-/-CL1^ Cas9)* and *Hira-* and *Fh1*-deficient cells (*Fh1^-/-CL1^ g1Hira)* and BioLuminiscence Imaging (BLI) and flux intensity at the experimental end point (week 9). **(c)** Representative H&A staining image of kidney capsule injected with *Fh1*-deficient cells (*Fh1^-/-CL1^ Cas9).* Scale bar represents 1mm. **(d)** Representation of luminescence signal by means of BioLuminiscence Imaging (BLI) and flux intensity normalised to time 0 of cells (*Fh1^fl/fl^ Cas9* and *Fh1^-/-CL1^ Cas9)* injected in the flanks of the mice (n= 10 tumours). **(e)** Representation of luminescence signal by means of BioLuminiscence Imaging (BLI) and flux intensity normalised to day 1 (day after surgery) of *Fh1^fl/fl^ Cas9* and *Fh1^-/-CL1^ Cas9* cells injected in the kidney capsule (n=4 kidneys). **(f)** qRT-PCR showing expression levels for *FH* and *HIRA* for two HLRCC patient samples and three adjacent normal tissue samples (n= 3 normal, n=2 HLRCC). RPLP0 was used as a housekeeping gene. Error bars represent standard error of the mean (S.E.M). Statistic tests performed: two-tailed Mann-Whitney U test (a, d, e). Numbers represent p-value for all comparisons. No statistics were performed in panel f due to n<3.

**Supplementary Figure 4.**
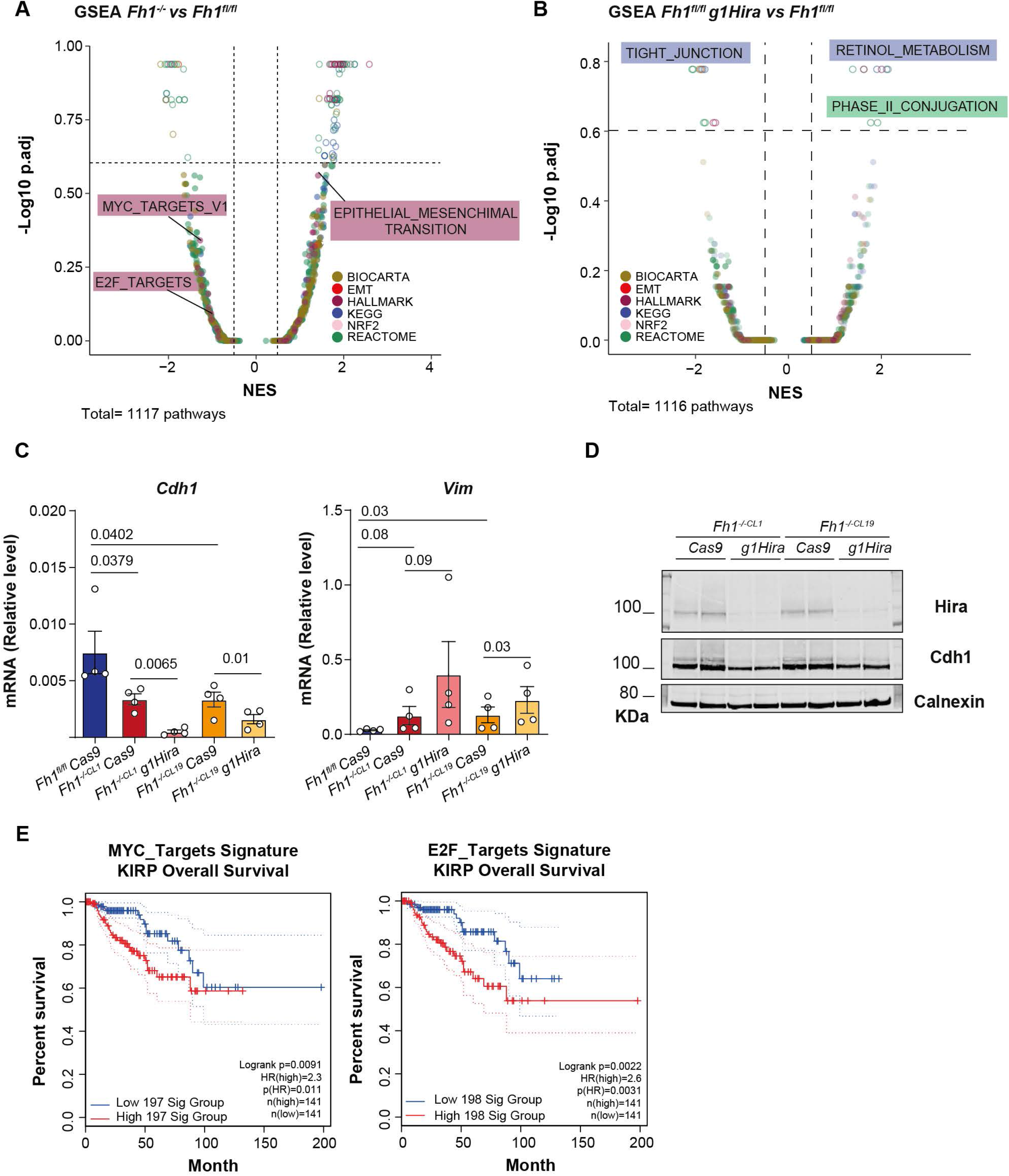
**(a**) Volcano plot representing the GSEA data from *Fh1^-/-^* vs *Fh1^fl/fl^.* **(b)** Volcano plot representing the GSEA from *Fh1^fl/fl^ g1Hira vs Fh1^fl/fl^*. **(c)** qRT-PCR showing expression levels for *Cdh1* and *Vim* in *Fh1*-proficient and deficient cells, and *Hira* and *Fh1*-deficient cells (n=4). β-Actin was used as a housekeeping gene. **(d)** Immunoblot of Hira and Cdh1 in *Fh1*-deficient cells and *Hira* and *Fh1*-deficient cells. Calnexin was use as a housekeeping gene. One representative experiment shown out of 3. **(e)** Percentage overall survival data associated to E2F targets and MYC targets signatures expression from Papillary type renal cancer (KIRP) using GEPIA(*22*). For volcano plots, signatures are highlighted depending on database represented. Error bars represent standard error of the mean (S.E.M). Statistic tests performed: two-tailed Student T test (c). Numbers represent p-value for all comparisons.

**Supplementary Figure 5.**
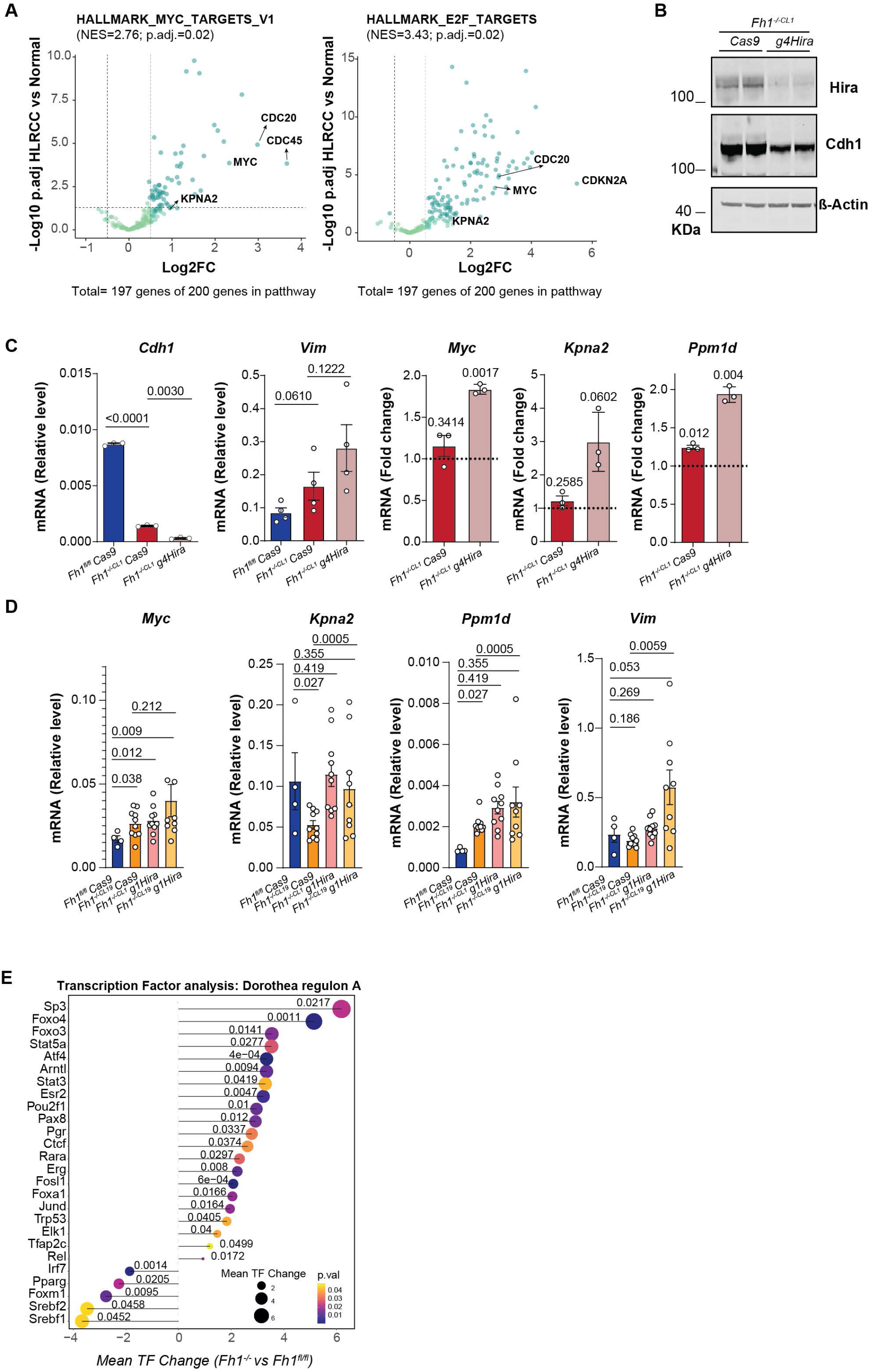
**(a)** Volcano plots of the genes present in the significantly upregulated signatures in the GSEA for HLRCC vs Normal (MYC_Targets_V1 and E2F_Targets). **(b)** Immunoblot of Hira and Cdh1 for Fh1-deficient cells (*Fh1^-/-CL1^*) and a second gRNA against *Hira* (*Fh1^-/-CL1^ g4Hira).* β-Actin was used as a housekeeping gene. One representative experiment shown out of 3. **(c)** qRT-PCR showing expression levels for *Cdh1, Vim, Myc, Kpna2* and *Ppm1d* of the previously mentioned cell lines. Dotted line represents control (*Fh1^fl/fl^ Cas9*). **(d)** qRT-PCR showing expression levels for *Myc, Kpna2, Ppm1d* and *Vim* in xenograft tumours generated with *Fh1*-proficient, *Fh1*-deficient, and *Hira*- and *Fh1*-deficient cells. **(e)** Lollipop chart with the transcription factor analysis from the RNA-seq data comparing *Fh1*-deficient cells versus *Fh1*-proficient cells. Error bars represent standard error of the mean (S.E.M). Statistic tests performed: two-tailed Student T test and one sample t test with a null hypothesis of 1 (c), two-tailed Mann-Whitney U test for unpaired comparisons and Wilcoxon test for paired comparisons (d). β-Actin used as a housekeeping gene. Numbers represent p-value for all comparisons.

**Supplementary Figure 6.**
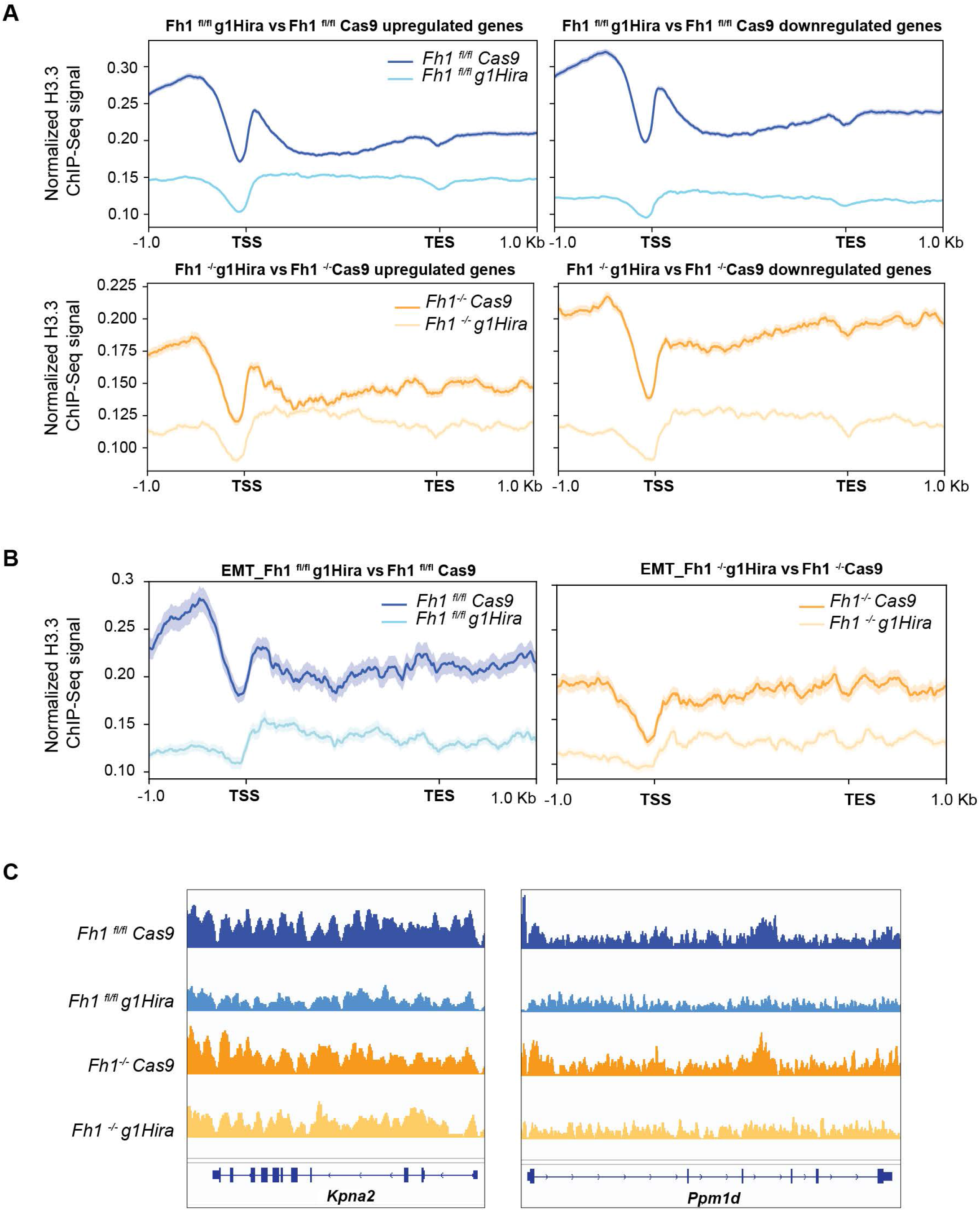
**(a)** Correlation of normalized H3.3 ChIP signal with upregulated and downregulated genes in the RNA-Seq results for the comparisons shown. **(b)** Normalized H3.3 ChIP signal associated with EMT signature expression for the comparisons shown. **(c)** IGV snapshot from H3.3 ChIP-Seq for the promoter regions of Kpna2 and Ppm1d genes. Shadows represent the S.E.M.

**Supplementary Figure 7.**
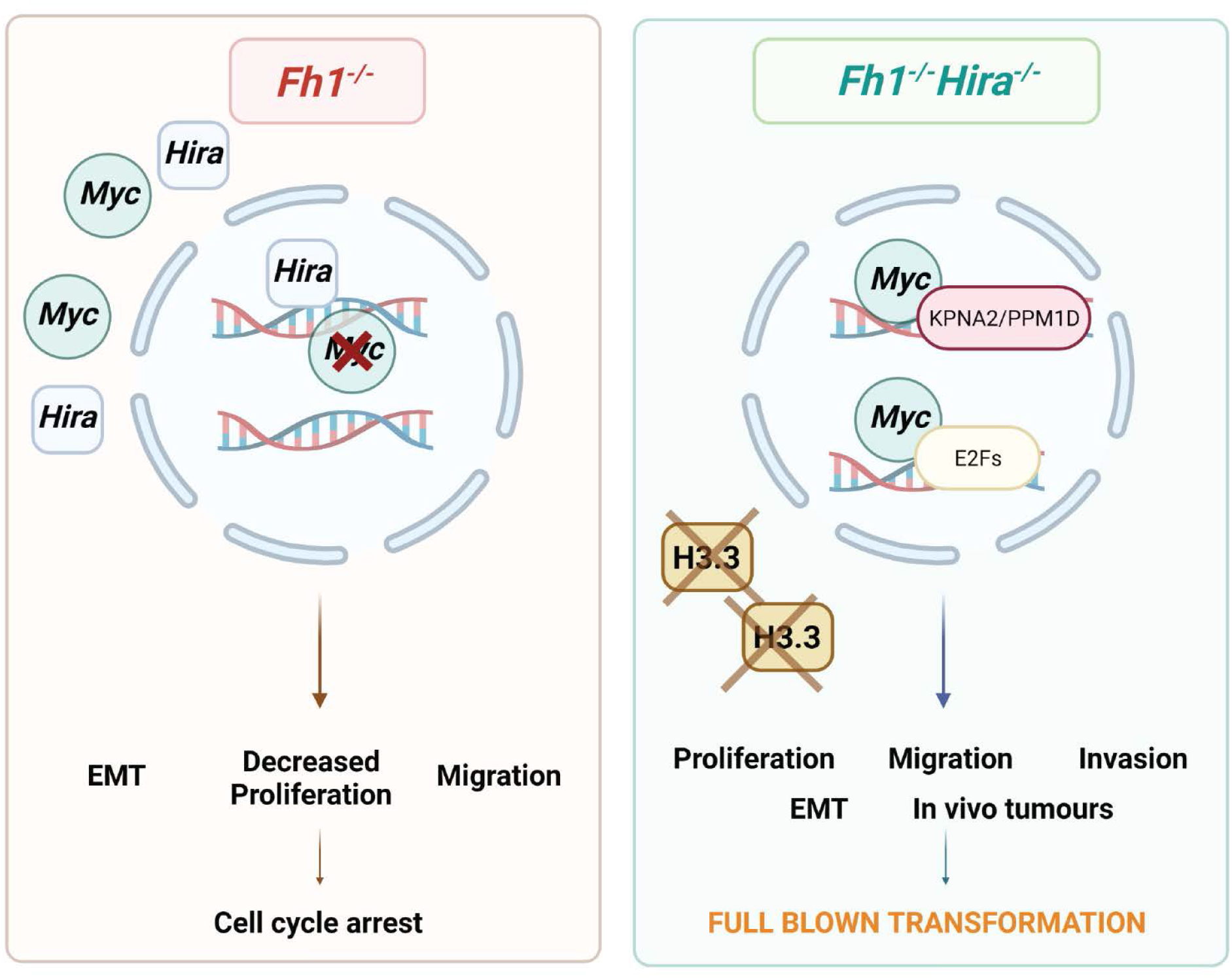
Schematic of the results obtained in this work. *Fh1* loss leads to a cell cycle arrest associated with increased migration and invasion. In this context, *Hira* blocks the accessibility of Myc into the nucleus, inhibiting the activation of its transcriptional programme, through a H3.3-independent mechanism. *Hira* loss in *Fh1*-deficient cells increases proliferation and enhances the invasive properties of the cells *in vitro* and *in vivo*. *Hira* loss in this context allows the binding of Myc in the chromatin and the activation of an oncogenic transcriptional programme allowing full blown transformation.

